# A short HBV RNA region induces RNR-R2 expression in non-cycling cells and in primary human hepatocytes

**DOI:** 10.1101/458679

**Authors:** Inna Ricardo-Lax, Karin Broennimann, Julia Adler, Eleftherios Michailidis, Ype P de Jong, Nina Reuven, Yosef Shaul

## Abstract

Hepatitis B virus infects non-dividing cells in which dNTPs are scarce. HBV replication requires dNTPs. To cope with this constraint the virus induces the DNA damage response (DDR) pathway culminating in RNR-R2 expression and the generation of an active RNR holoenzyme, the key regulator of dNTP levels. Previously we reported that the HBx open reading frame (ORF) triggers this pathway. Unexpectedly however, we report here that the production of HBx protein is not essential. We found that a small region of 125 bases within the HBx transcript is sufficient to induce RNR-R2 expression in growth arrested HepG2 cells and in primary human hepatocytes (PHH). The observed HBx embedded regulatory element is named ERE. We demonstrate that ERE is functional as a positive strand RNA polymerase-II transcript. Remarkably, ERE is sufficient to induce the Chk1-E2F1-RNR-R2 DDR pathway, previously reported to be activated by HBV. Furthermore, we found that ERE activates ATR but not ATM in eliciting this DDR pathway in upregulating RNR-R2. These findings demonstrate the multitasking role of HBV transcripts in mediating virus-host cell interaction, a mechanism that explains how such a small genome effectively serves such a pervasive virus.

**Author summary:** The hepatitis B virus (HBV) infects the human liver and over 250 million people worldwide are chronically infected with HBV and at risk for cirrhosis and liver cancer. HBV has a very small DNA genome with only four genes, much fewer than other viruses. For propagation the virus consumes dNTPs, the building blocks of DNA, in much higher amounts than the infected cells provide. To cope with this constraint, the virus manipulates the cells to increase the production of dNTPs. We found that the virus activates the cellular response to DNA damage upon which the cells increase the production of dNTPs, but instead of repairing cellular DNA, the virus uses them for production of its own DNA. Usually viruses manipulate host cells with one or more of their unique proteins, however the small HBV genome cannot afford having such a unique gene and protein. Instead, we found that here the virus relies on RNA to manipulate the host cells. Our findings highlight the unprecedented principle of a multitasking viral RNA that is not only designed to be translated into proteins but also harbors an independent role in activating the cellular DNA damage response.

## Introduction

Hepatitis B virus (HBV) is a non-cytopathic enveloped virus containing a small circular partially double-stranded DNA genome. Upon entering the cell the genome is converted into a covalently closed circular DNA (cccDNA), the viral transcription template [1]. The HBV genome harbors enhancers and promoters regulating the transcription of a number of positive strand transcripts. The generated RNA species are nuclear exported by a unique mechanism that is not entirely understood but requires a RNA region shared by all the viral mRNAs called PRE (posttranscriptional regulatory element) [2,3]. An exceptional RNA species is the longest viral transcript that accumulates in the nucleus [4].

HBV, a hepatotropic virus, infects hepatocytes, where the virus successfully propagates and produces infectious progeny. Chronic HBV infection gives rise to a wide range of clinical manifestations extending from acute and chronic hepatitis to hepatocellular carcinoma (HCC). The HBV-hepatocyte tropism is determined by both receptor and postreceptor mechanisms. The latter is mediated by the viral cccDNA enhancers recruiting liver enriched transcription factors to initiate transcription of the viral genome [5]. The HBV receptor tropism is determined by a liver-specific bile acid transporter, the sodium taurocholate co-transporting polypeptide (NTCP) [6].

There is a certain level of similarity between the life cycle of HIV and HBV. Both infect non-cycling cells and replicate via reverse transcription, mediated by a reverse transcriptase (RTase), of the pre-genomic RNA. However, unlike the HIV undergoing reverse-transcription early upon infection [7], HBV utilizes RTase at the final stage of its life cycle in the process of progeny maturation. Whereas HIV RTase activity is dispensable after the first round of DNA synthesis, HBV RTase is active as long as new viruses are formed. As the result, HBV consumes much larger amounts of deoxynucleotides (dNTPs), the building blocks of DNA, than HIV. The dNTP pool is extremely low in non-cycling cells, which leads to the question how the HBV RTase is active in DNA synthesis under this condition.

In cycling cells, the ribonucleotide reductase (RNR) holoenzyme catalyzes the synthesis of dNTPs from rNTPs. The RNR complex contains large R1 and small R2 subunits. The R2 subunit (RNR-R2) is encoded by the *RRM2* gene that is exclusively expressed at the entry to S phase of cycling cells. In non-cycling cells, the RNR enzyme is not functional, further reiterating the question of how HBV DNA is synthesized under this condition. Furthermore, hydroxyurea (HU), a specific RNR inhibitor blocks HBV production [8], demonstrating the necessity of the RNR activity for the HBV life cycle. Upregulation of RNR-R2 expression by HBV in non-cycling cells [8,9] is a critical step in ensuring productive infection.

A well-known mechanism of RNR-R2 upregulation is via cell proliferation pathways. Over-expression of HBx in rat hepatocytes induces some of the cell proliferation hallmarks [10] and HBV infection of PHH deregulates cell cycle to increase the G2/M population [11]. However, the role of HBV, in particular the HBx protein, in inducing proliferation of non-cycling cells is highly controversial [12]. On the other hand, there is an alternative pathway for RNR-R2 expression upregulation triggered by DNA damage response (DDR) [13,14]. Under this condition the E2F1 transcription factor, a key regulator of RNR-R2 transcription, is phosphorylated by the Chk1 S/T kinase [15] a process that potentiates its activity. We previously reported that HBV induces RNR-R2 via the Chk1-E2F1 pathway [9] (Fig S1 and S2). Here we investigated the HBV element that induces RNR-R2 expression. We found that the HBx gene is sufficient to induce RNR-R2 but unexpectedly, the intact HBx ORF is not required for this process. A small region of HBV RNA within the HBx gene is enough. Thus, the compact HBV genome with a low number of ORFs has adapted new mechanisms of regulating virus-host cell interaction by acquiring a novel mRNA moonlighting activity.

## Results

### Of the HBV ORFs, only HBx induced RNR-R2 expression

The HBx ORF in isolation under either a CMV or Adenovirus promoter is sufficient to induce RNR-R2 expression [8,10]. We transduced cells with a lentivector (LV) containing either the 1.3xHBV genome or the HBx gene in isolation. We revealed that HBx was as effective as HBV in upregulating RNR-R2 expression (Fig S2 A - C). To demonstrate that in the context of the intact HBV genome the HBx ORF was responsible for the RNR-R2 upregulation, we inserted a stop codon to construct the 1.3xHBV HBx G27-stop mutant (Fig 1A). Both HBx copies in the 1.3xHBx plasmid were mutated but the overlapping Pol ORF was preserved. This stop codon successfully eliminates HBx production (see below). Unexpectedly, we found that this mutant was active in upregulating RNR-R2 expression at the protein (Fig 1B) and RNA levels (Fig 1C) in noncycling HepG2 cells. These data suggest either that an intact HBx ORF is not required for RNR-R2 upregulation or that the other HBV ORFs are involved in this process as well. To rule out the latter possibility we replaced the HBsAg or HBc ORF with a GFP ORF (Fig. 1A). The obtained data show that the HBsAg knockout mutant was active in induction of RNR-R2 protein (Fig 1B) and RNA (Fig 1C) as was the HBc knockout mutant (Fig. 1D (Protein) and 1E (RNA)). Given the compact structure of the HBV genome, the Pol ORF was hampered in the context of these mutants as well, therefore Pol ORF is unlikely to be involved in the RNR-R2 induction. These results suggest that neither of the other HBV ORFs is sufficient in inducing RNR-R2 upregulation in quiescent cells.

**Fig 1.**
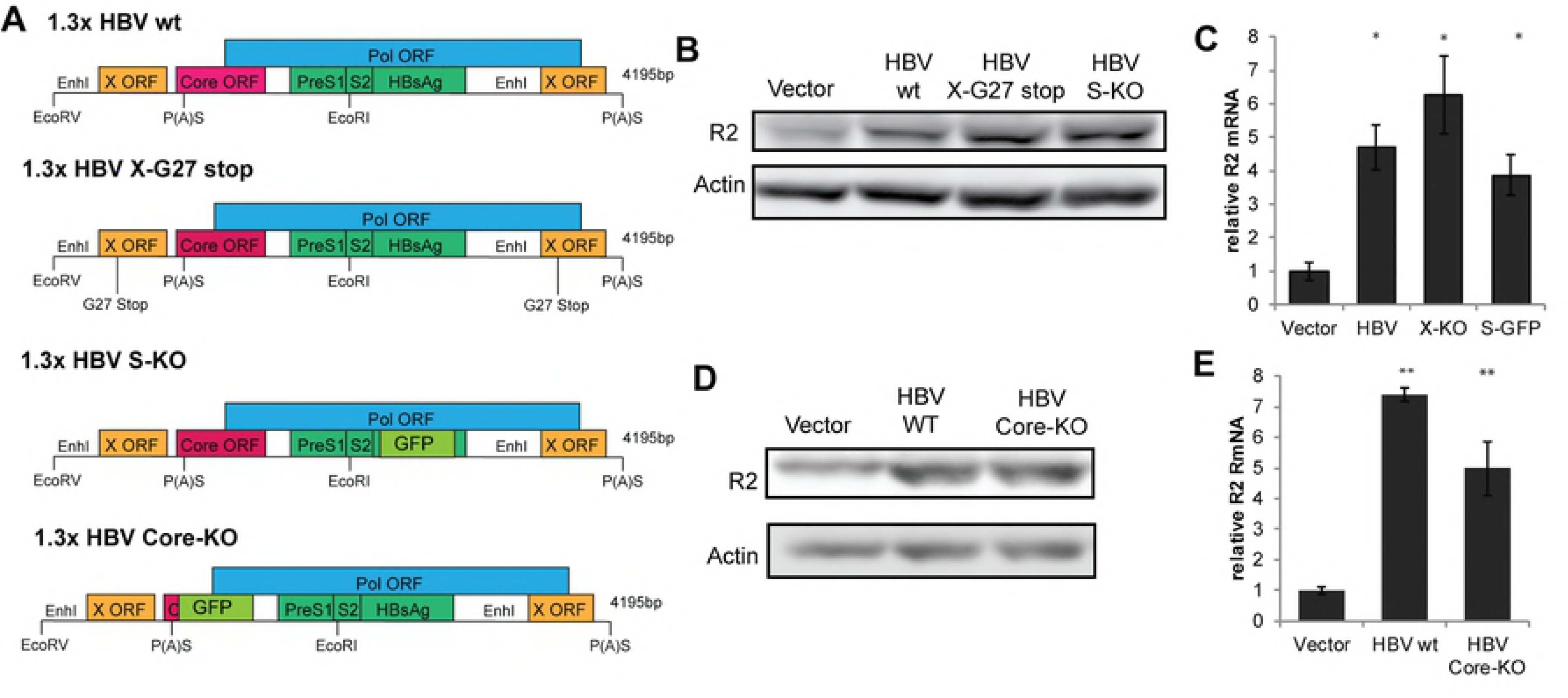
None of the HBV ORFs is required for RNR-R2 upregulation in non-cycling cells. A) Schematic depiction of the 1.3xHBV genome organization with the major ORFs (top) and with the sites of the insertion of the stop codon in the HBx ORF (bottom). B) The level of RNR-R2 protein after transduction of non-cycling HepG2 with the X-G27 and S-KO constructs described in A. Actin was used as loading control. C) As in B but the level of R2 transcripts measured by qRT-PCR is shown. *-p-value< 0.05 was calculated for each sample compared to the vector control using Student’s t-test. D) The level of RNR-R2 protein after the transduction of non-cycling HepG2 with the Core-KO construct depicted in A. E) As in D, but the level of R2 transcripts measured by qRT-PCR is shown. **-p-value < 0.01 was calculated for each sample compared to the vector control.

### The HBx sequence but not an intact ORF is required for RNR-R2 induction

Since the HBx ORF in isolation and the 1.3xHBV HBx G27stop mutant both induced RNR-R2 expression, we reasoned that a region in HBx but not the intact HBx ORF regulates this process. To address this possibility we introduced the stop codon in the HBx ORF in isolation. The plasmids contain a HA-tag for protein detection and the endogenous 3’UTR of HBx (Fig. 2A). Remarkably, the G27stop mutant was active in RNR-R2 induction under this condition (Fig 2B). The HBx ORF contains two additional internal ATG residues at the positions of amino acids 79 and 103 that might initiate translation. We consequently mutated the next codon to a stop codon and inserted a stop codon immediately downstream of the HA-tag. Interestingly, all these mutants were fully active in RNR-R2 induction (Fig. 2B). These constructs were all efficiently expressed at the mRNA level (Fig. 2C), but as expected, only the WT construct expressed the HA-HBx protein (Fig 2D).

**Fig 2.**
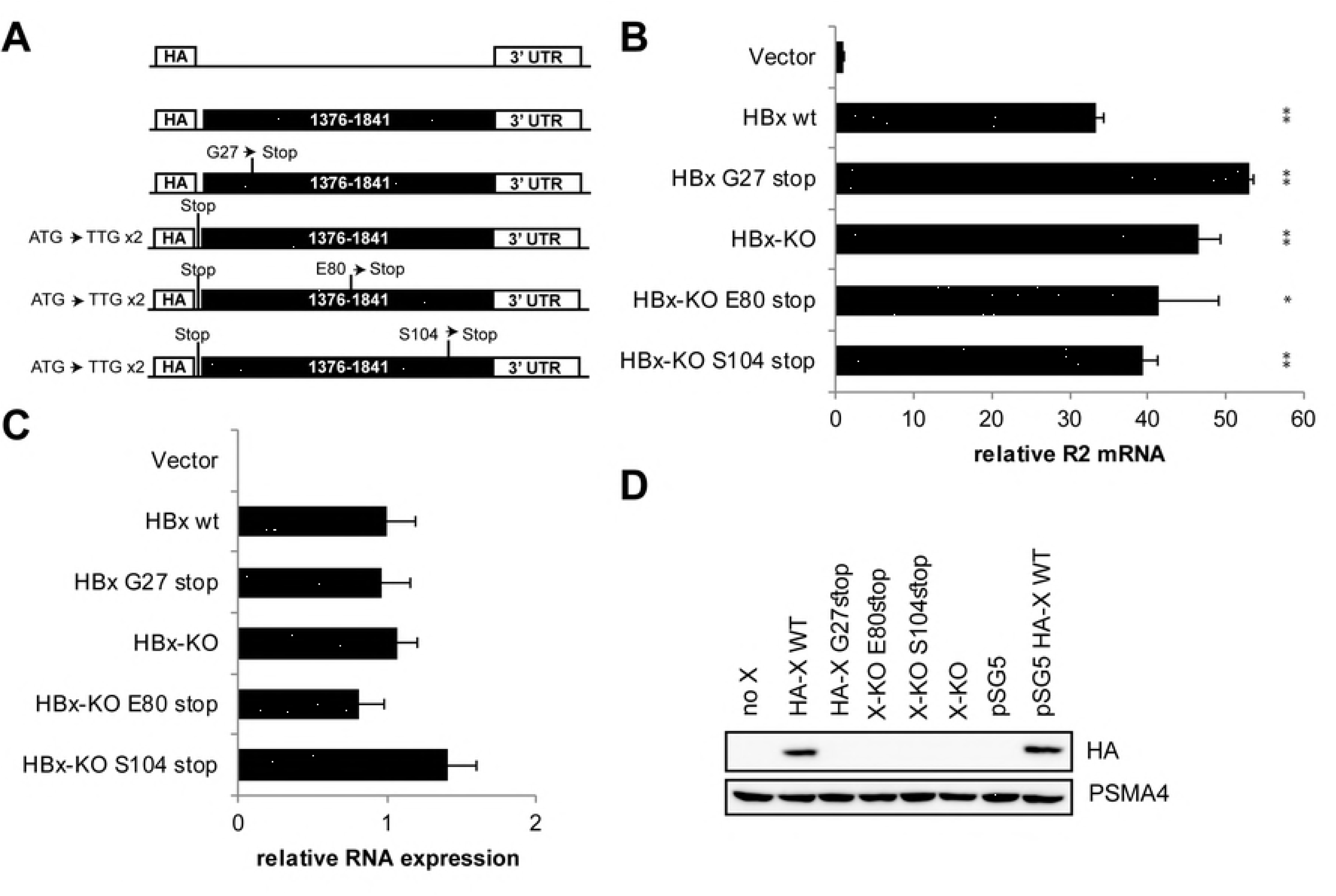
Translation of HBx ORF is not required for RNR-R2 upregulation. A) Schematic depiction of the HA-HBx ORF (black regions) with the inserted mutations, all cloned in a lenti-vector (LV). B, C) non-cycling HepG2 cells were transduced with the HBx gene mutants of panel A and relative level of the expressed R2 RNA (B) and Lentiviral construct expression (C) were quantified by qRT-PCR. *p-value<0.05, **p-value<0.01 were calculated for each sample compared to the vector control using Student’s t-test D) The constructs in A were transfected into HEK293 cells. HA-HBx protein levels were measured by western blot with anti-HA antibody PSMA4 was used as a loading control. The pSG5-empty and pSG5-HA-HBx constructs [40] were used as negative and positive controls respectively.

Since the Pol ORF partially overlaps with the HBx ORF, we also introduced a stop codon after the HA-sequence in the Pol reading frame (Fig. S3). This mutant was also active in RNR-R2 induction, further ruling out Pol ORF involvement. These data suggest that the HBx gene region, but not the protein, is required and sufficient for RNR-R2 induction in non-cycling HepG2 cells.

### Delineation of the minimal active HBx ORF region

Having demonstrated that an intact HBx ORF is dispensable for RNR-R2 upregulation, we next deleted the 3’ UTR region and found that this deletion did not compromise RNR-R2 activation. This was also the case with the deletion of the 5’ sequences including the HA-tag (Fig S4). Finally we deleted the entire HBx coding region, and retained only the HA sequence followed immediately by the 3’ UTR (no X in Fig 3A). This construct was inactive in RNR-R2 induction (Fig 3B) despite being efficiently expressed, suggesting that the HBx coding region, but not the flanking 5’ or 3’ sequences are responsible for RNR-R2 upregulation.

**Fig 3.**
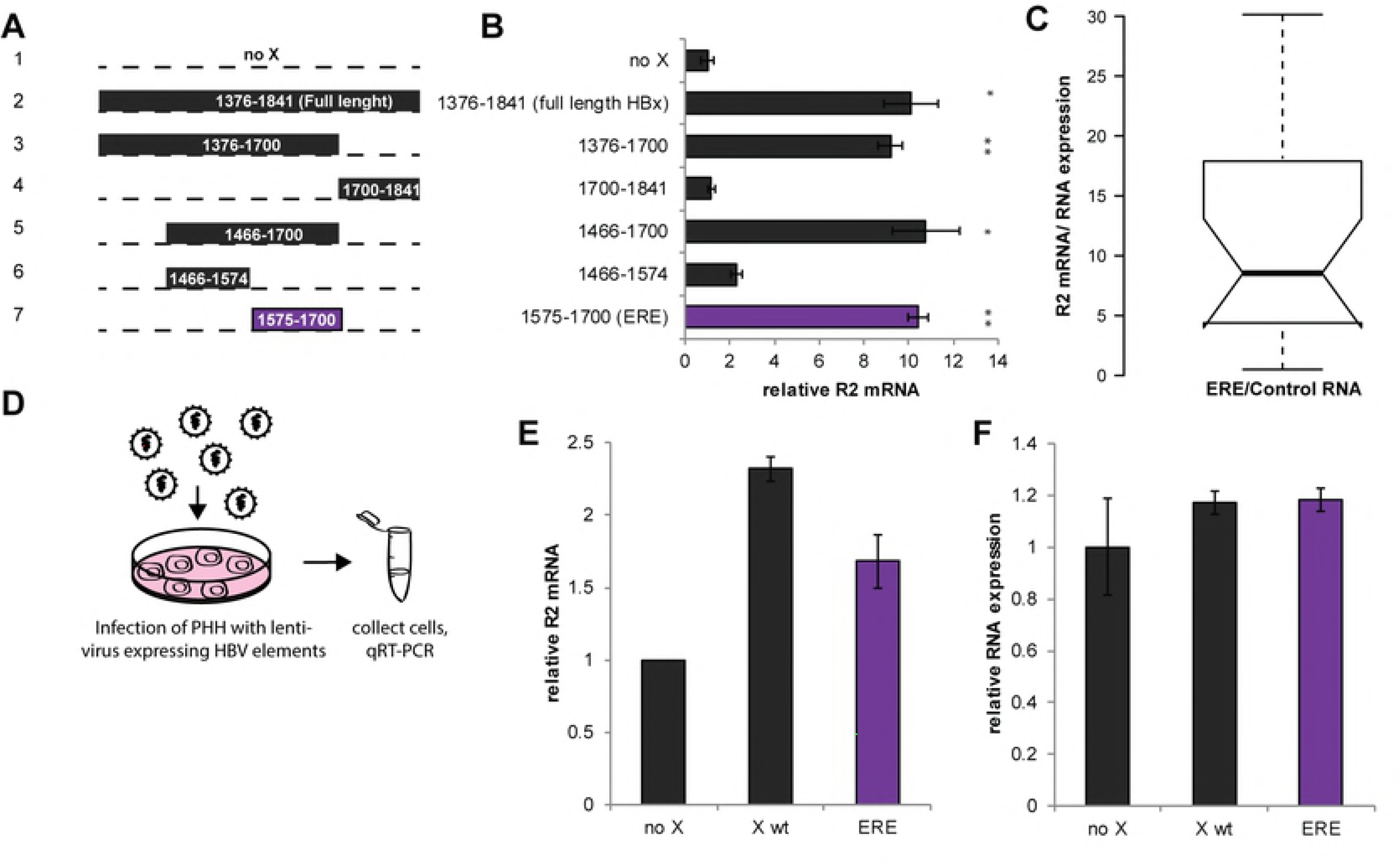
Delineation of the HBx RNA minimal sequence required for RNR-R2 upregulation. A) Schematic representation of the truncated HBx gene in LV constructs. The black regions were cloned for expression. The standard numbering from the EcoRI site in the HBV genome is indicated. B) The constructs in panel A were transduced into non-cycling HepG2 cells. Relative RNR-R2 mRNA levels were measured by qRT-PCR from three biological replicates. *-p-value<0.05, **-p-value<0.01 were calculated for each sample compared to vector control using Student’s t-test. C) A boxplot of the fold changes of RNR-R2 expression between ERE and the control RNA of 23 biological replicates is shown. The R2 level was normalized to the level of expression of the HBx fragments. Student’s T-test was performed and the results are highly significant (p < 0.0001). D) Primary human hepatocytes were transduced with the lentiviral constructs number 1, 2 and 7 in panel A. E) Following transduction in D, relative RNR-R2 mRNA levels of biological duplicates were measured by qRT-PCR. F) As in E, LV RNA levels were measured by qRT-PCR to assure similar expression of the constructs.

Next, we constructed a series of truncation mutants, to delineate the HBx RNA region that induces RNR-R2 expression (Fig 3A). Remarkably, a 125 bases long fragment corresponding to nucleotides 1575–1700 (taking the unique HBV EcoRI site as 1) was sufficient for RNR-R2 induction to the level obtained by the full length HBx ORF (Fig 3B). We refer to this positive RNA fragment as embedded regulatory element (ERE). The immediate ERE upstream sequence (nucleotides number 1466–1574) was marginally active and was used as a negative control RNA in following experiments. To get a more statistical view we repeated the experiment (23 biological repeats) and found that on average the ERE fragment leads to about 8 folds higher RNR-R2 expression levels than the negative control RNA (Fig 3C).

We have previously reported that HBV induces RNR-R2 over two folds in infected primary human hepatocytes (PHH) [9]. To show whether ERE is active under physiological conditions, we transduced PHH and revealed that WT HBx increases the level of the RNR-R2 expression (Fig 3D-F), close to the level observed after HBV infection. ERE was active in inducing RNR-R2 expression as well. These data suggest that ERE, a small region of the HBx ORF is responsible for the RNR-R2 upregulation both in non-cycling HepG2 cells and in primary human hepatocytes.

### ERE is active as a positive strand Pol-II

To validate the observation that the RNR-R2 upregulation is mediated by transcribed ERE and not by the DNA itself, we cloned ERE with or without an upstream CMV promoter. Our analysis revealed that RNR-R2 was induced more efficiently in the presence of the CMV promoter (Fig 4A), suggesting that the active molecule is the RNA. To test whether the ERE is active in an orientation-dependent manner, the ERE fragment was cloned under the same regulatory elements but in the opposite orientation (Fig 4B). RNR-R2 was only induced by the positive orientation construct (Fig 4C) even though the expression of the reverse construct was high (Fig 4D). These results suggest that a plus-strand RNA sequence is responsible for the RNR-R2 induction.

**Fig 4.**
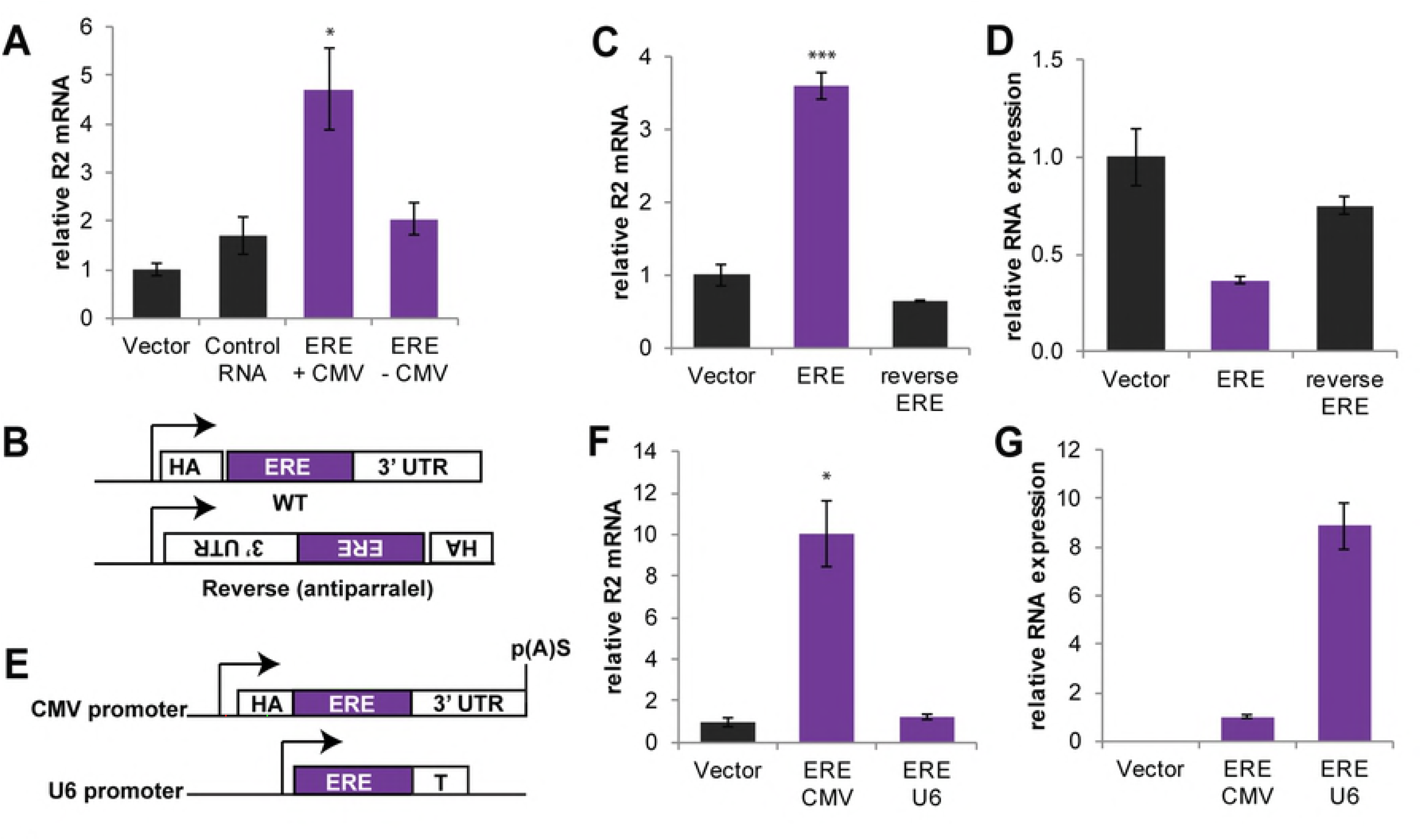
Functional ERE is a sense Pol-II transcript. A) The ERE fragment was cloned into a lentiviral construct with (+) or without (-) a CMV promoter. An empty lentivector and the control RNA fragment under CMV promoter were used as negative controls. Non-cycling HepG2 cells were transduced with the indicated lentiviral constructs. Shown are relative RNR-R2 mRNA levels, measured by qRT-PCR from three biological replicates. Student’s t-test was performed comparing all samples to the vector control, only ERE + CMV was significantly (*-p<0.05) changed. B) The ERE expression unit was cloned into lentivectors in sense and anti-sense orientation. C) Non-cycling HepG2 cells were transduced with constructs depicted in B. Relative RNR-R2 mRNA levels were measured by qRT-PCR from three biological replicates. T-test was performed comparing the samples with the vector control, only sense ERE changed significantly (***-p<0.001) D) Viral RNA expression levels were measured by qRT-PCR. E) The ERE under Pol-II expression using a CMV promoter and under Pol-III expression using a U6 promoter and specific transcription termination (depicted as T in the illustration) were constructed in the LV backbone. F) The constructs in D were transduced into non-cycling HepG2 cells. RNR-R2 mRNA levels were measured by qRT-PCR. As a control, we used the same CMV-based vector, lacking the ERE sequence. T-test was performed comparing the samples with the vector control, only CMV ERE changed significantly (*-p<0.05) G) ERE expression levels were measured by qRT-PCR.

Next, we asked whether ERE expression by a RNA polymerase (Pol)-III promoter is functional. To this end we cloned the ERE fragment under the Pol-III U6 promoter (Fig 4E), which is often utilized for expression of small regulatory RNAs. For Pol-II control, we used the original construct, bearing a CMV promoter, efficiently targeted by Pol-II to express mRNAs. Interestingly, RNR-R2 was induced only when the ERE fragment was expressed from a Pol-II promoter, but not from a Pol-III promoter (Fig 4F), even though ERE expression was about eight fold higher under the U6 promoter (Fig 4G). This suggests that the ERE fragment is only active when expressed like the native HBV transcripts.

### ERE induces the Chk1-E2F1-RNR-R2 axis

Previously, we reported that HBV induces RNR-R2 in quiescent cells by activating the DNA damage response [9]. Lenti-HBV transduction leads to an increase in Chk1 phosphorylation, which is a marker for its activation, and Chk1 kinase activity is required for RNR-R2 induction in non-cycling cells. To determine whether ERE is sufficient to increase Chk1 S345 phosphorylation, we transduced HepG2 cells and assayed for Chk1 protein levels and phosphorylation. Calf intestinal phosphatase (CIP) was used to validate phosphorylation. As expected, the positive control HBx ORF induced Chk1 phosphorylation and accumulation (Fig 5A). Remarkably, expression of ERE in isolation was sufficient to induce Chk1 activity.

**Fig 5.**
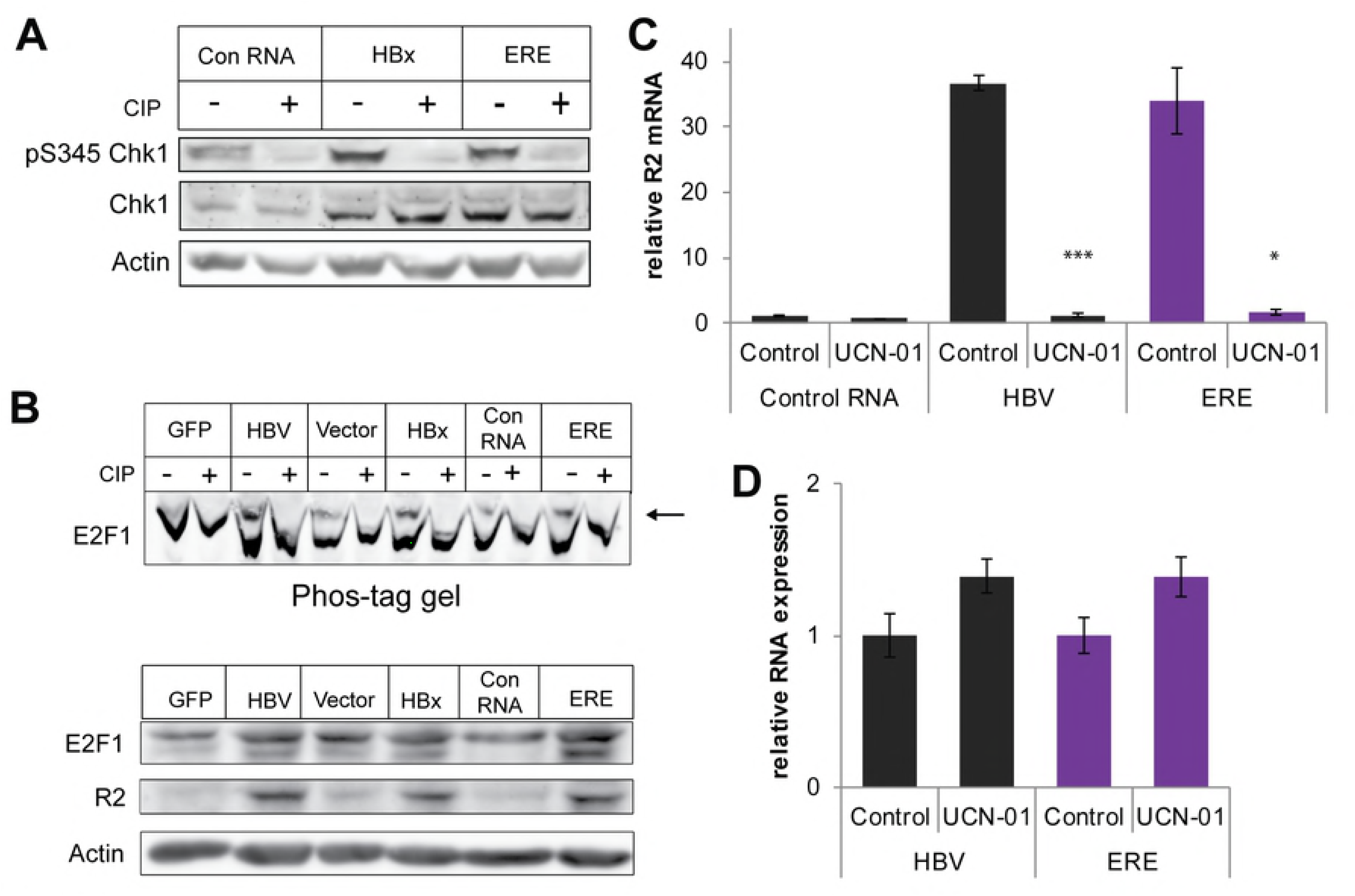
ERE activates DNA damage response. A) HepG2 cells were transduced with LV containing the indicated constructs. Phospho-S345 Chk1 levels were measured by western blot, using a specific antibody, and the phosphorylated nature of the bands was validated by Calf Intestinal Phosphatase (CIP) treatment. Total Chk1 protein levels were measured as well. B) Upper panel: Non-cycling HepG2 cells were transduced with the indicated LV constructs. The protein samples were treated with CIP or left untreated, and subjected to Phos-Tag SDS-PAGE gel, to separate the phosphorylated E2F1 forms. The arrow marks the phospho-E2F1 band. Lower panel: Extracts from the indicated transduced constructs were analyzed by western blot for E2F1 and RNR-R2 proteins. C) Non-cycling HepG2 cells were transduced with the indicated constructs and treated with 1μM UCN-01, an inhibitor of Chk1 kinase activity, for 24h. Relative RNR-R2 mRNA levels were measured by qRT-PCR from three biological replicates. A Students T-test was performed comparing the UCN treated and untreated samples. *-p-value<0.05, ***-p-value<0.001 D) As in E, but the level of the RNA expressed from the transduced constructs was measured.

The pS345 Chk1 phosphorylates E2F1, the transcription activator of the *RRM2* gene. We measured the levels of phospho-E2F1 with a phos-tag gel in the presence and absence of ERE. As expected, under HBV expression, E2F1 was markedly phosphorylated. Similar results were obtained with ERE expression. A regular SDS-PAGE gel showed that total E2F1 protein levels were increased with ERE. These results suggest that ERE expression leads to E2F1 phosphorylation and upregulation. RNR-R2 protein levels were also measured, and were markedly induced with Lenti-HBV, Lenti-HBx and the ERE constructs (Fig 5B).

Next, we wished to determine whether Chk1 kinase activity is required for RNR-R2 induction by ERE. We used a specific Chk1 inhibitor UCN-01 and found that UCN-01 completely prevented HBV induced RNR-R2 expression (Fig 5C), in agreement with our previous report [9]. Interestingly under this condition RNR-R2 upregulation by ERE was also inhibited. We measured ERE and HBV relative RNA levels to rule out the possibility that UCN-01 reduced ERE expression (Fig 5D). These data suggest that ERE is sufficient to induce the Chk1-E2F1-*RRM2* DDR axis.

### ERE activates DDR via ATR and not ATM

The main kinase responsible for Chk1 S345 phosphorylation and activation is ATR [16–18]. We tested the possibility of ATR involvement in ERE induced DDR. We used Caffeine, an inhibitor of both ATM and ATR, and followed RNR-R2 mRNA induction (Fig 6A, Fig S5A). Caffeine significantly inhibited RNR-R2 induction, despite the fact that ERE was well expressed (Fig 6B, Fig S5B). We also treated the cells with a specific ATM inhibitor, KU55933, and found that RNR-R2 induction was unaffected by this treatment (Fig S5A). X-ray irradiation induces DNA damage, ATM activation and p21, a known target of ATM, is upregulated. Indeed KU55933 reduced p21 induction upon ionizing irradiation (Fig S5C), providing a positive control for the activity of the ATM inhibitor in our system. These data suggest that ERE did not activate ATM. In contrast, when we used a specific ATR inhibitor, AZD6738, RNR-R2 induction by ERE was significantly reduced (Fig 6A), in spite the fact that ERE was well expressed (Fig 6B). Next, we measured ATR phosphorylation of Thr1989, which is a marker for ATR activation [19,20] in the presence of ERE and revealed that the level of phosphorylated ATR, but not total ATR was increased (Fig 6C and D). These data support the model that HBV ERE induces RNR-R2 expression by activating the ATR-Chk1-E2F1 axis of DDR (Fig 6E).

**Fig6.**
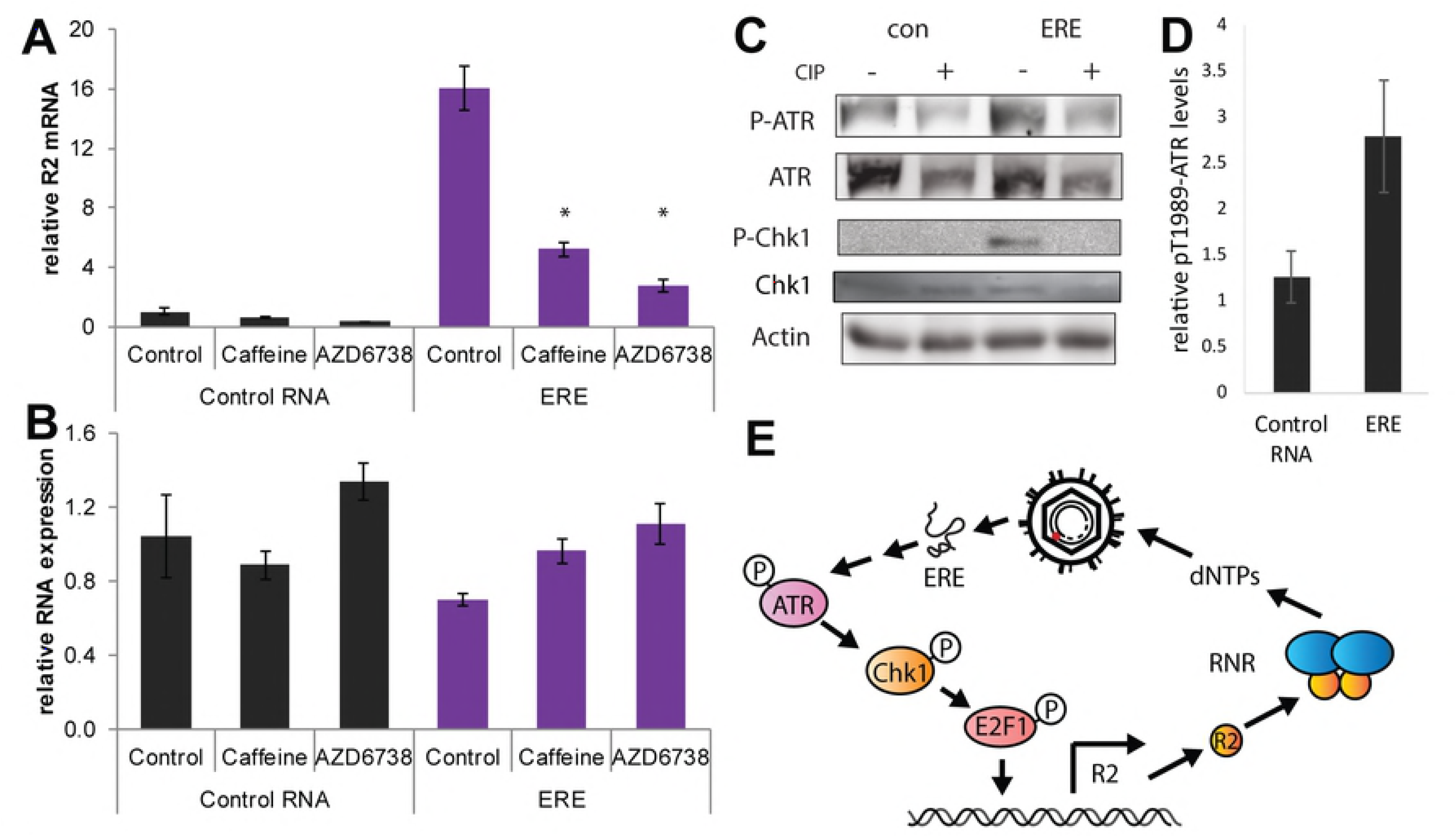
ERE activates ATR. A) Non-cycling HepG2 cells were transduced with LV-constructs containing ERE or a negative control RNA. After 48h, cells were treated with 2μM Caffeine or 10μM AZD6738, a specific ATR inhibitor, or left untreated. RNR-R2 mRNA levels were measured 24h after treatment with qRT-PCR from three biological replicates. A Students T-test was performed comparing the control of each sample to the treatments. *-p-value<0.05 B) The samples in A were analyzed for lentiviral RNA expression using qRT-PCR to validate similar expression levels. C) PhosphoT1989 ATR, PhosphoS345 Chk1, Chk1 and total ATR levels were examined in the presence and absence of ERE by western blot. CIP was used to evaluate the specificity of the phospho-antibodies. Actin was used as loading control. D) Quantification of P-ATR levels of five biological replicates of C). E) The proposed signaling pathway activated by ERE.

## Discussion

We investigate the molecular basis of HBV-host cell interaction after infection. This is a challenging event particularly for a virus with such a small DNA genome encoding just a few ORFs. HBV exploits cellular mechanisms for almost every step in its life cycle. Here we addressed the question of how the virus ensures optimal dNTP levels for DNA synthesis of the viral progeny leading to productive infection. Since new HBV virions are generated in non-dividing cells, the question of what is the source of the dNTPs, the DNA building blocks, is very important. Initially we observed that the HBV-positive HepG2-2-15 cells produce large amounts of thymidine [8] indicating a role of HBV in biosynthesis of dNTPs. Follow-up studies revealed that HBx induces RNR-R2 expression in non-cycling cells. Since in our experimental system HBx does not induce cell proliferation we looked for an alternative pathway that upregulates RNR-R2 expression. Indeed, we demonstrated that the Chk1-E2F1-RNR-R2 DDR circuit is activated by HBV and HBx [9].

Based on detailed mutagenesis studies we report here that the integrity of the HBx ORF is dispensable and that RNR-R2 can be activated by a small 125 bases long RNA sequence inside the HBx ORF that we named embedded regulatory element, or ERE in short. ERE is functional as a positive-strand RNA and therefore all the HBV transcripts contain ERE. Given the finding that ERE is active when transcribed by Pol-II, it is very unlikely that ERE is transcribed in isolation by a novel transcript that so far has escaped detection.

ERE activates the ATR-Chk1-E2F-RNR-R2 DDR axis, like HBV, and therefore we believe that ERE activity is the only elicitor of the RNR-R2 activation by the virus. Indeed the fold of activation obtained by ERE in isolation is similar to that obtained either by the HBV 1.3xDNA genome or by the HBx ORF. However, we cannot experimentally rule out the possibility that some other viral components are involved because of the compact HBV genome that does not allow any significant mutation such as ERE deletion.

Activation of the different DDR pathways appears to play other important roles in the HBV life cycle. The formation of the episomal HBV cccDNA from rcDNA relies on cellular enzymes that are components of a DDR pathway sensing nicked DNA to be repaired [21]. It is very likely that this process also facilitates HBV DNA integration, a process that has an important implication in virus mediated pathogenesis [22]. Unlike the ERE that is actively involved in the activation of the described pathway of RNR-R2 activation, the DDR involved in cccDNA formation appears to be a more passive and default process, stimulated by the DNA nick.

There is accumulating evidence that other DNA viruses activate some of the cellular DDR pathways upon infection. Of particular interest are the human papillomaviruses (HPV). HPV induces breaks into the viral DNA in the process of genome amplification [23]. Like with HBV, HPV31 positive cells have an increased dNTP pool, which is in direct correlation with the RNR-R2 levels [24]. In that case, also the ATR-Chk1-E2F1 pathway is activated. However, unlike with HBV, in the case of HPV31 this pathway is activated by the viral E7 protein and not by an RNA element, although this possibility was not investigated. The Epstein-Barr virus (EBV) also has been reported to elicit a DDR pathway during primary infection and lytic reactivation to complete the replication process of viral DNA [25]. This has also been demonstrated in the case of Kaposi’s sarcoma-associated herpesvirus (KSHV) [26]. On the other hand the human adenovirus type 5 (Ad5) targets the cellular Mre11-Rad50-Nbs1 complex (MRN) in inhibiting DDR [27,28]. The emerging notion is that many DNA viruses target cellular DDR pathways to serve their different needs. In the reported cases, viral proteins mediate the DDR pathway manipulations. HBV is the first case where a region of RNA induces the cellular DDR. The 3.2kbp genome of HBV is very small with a very limited number of ORFs of which only HBx is not a structural protein with certain regulatory functions. The question is how such a limited number of genes is sufficient in exploiting the cellular pathways and recruiting essential cellular machineries in programming productive infection. It appears that HBV transcripts rather than proteins fulfill these tasks. The HBV transcripts are made of a number of functional entities embedded inside the RNA sequence. These include the encapsidation signal necessary for pgRNA packaging [29,30], the PRE regulating the nuclear export of the HBV transcripts [2,3,31], the RNA region involved in destabilizing HBV RNA in response to cytokines treatments [32,33] and the ERE described here. It is therefore possible that with the minimization of the HBV genome, some of the essential activities have become adapted to be executed by the viral RNA and therefore the emergence of multitasking HBV RNA is an important evolutionary step in establishing the hepadnavirus family.

## Methods

### Tissue culture, treatments and reagents

HepG2, HEK293 and HEK293T (ATCC) cells were cultured in Dulbecco’s modified Eagle’s medium (Gibco) supplemented with 8% fetal bovine serum (Gibco) and 100 U/ml of penicillin and 100 μg/ml of streptomycin (Biological Industries). To obtain quiescent HepG2, the medium was supplemented with 2% dimethyl sulfoxide (DMSO) (Sigma-Aldrich) for at least six days.

The reagents used were UCN-01 (7-hydroxystaurosporine) and Caffeine (Sigma), KU55933 and AZD6738 (ApexBio).

For X-Ray, a XRAD 320 by Precision X-Ray was used.

### Preparation of lentiviral transducing particles and transduction

Lentiviruses were produced as described [8] using the calcium phosphate method to transfect HEK293T. Lentivirion containing medium was filtered through 0.45μM cellulose acetate filter, and supplemented with 8μg/ml polybrene.

Medium was removed from the HepG2 cells and virion-containing medium was used to transduce the cells. 12–24h after infection the cells were washed five times in phosphate buffered saline (PBS) and fresh medium was added to the cells. Transduced cells were harvested after 72h.

### Primary human hepatocytes

For lentivirus production, 293T-LentiX cells were plated on poly-L lysine coated plates. The next day, cells were co-transfected with specific pLenti4 plasmids, together with VSV-G and Gag-Pol expressing packaging plasmids, using PEI transfection reagent. Lentivirus-containing medium was collected after 48 h, and cleared by filtering through a 0.45μM low protein-binding filter and centrifugation at 4,000 rpm for 10 min. Virus was concentrated by ultracentrifugation using a SW41 Ti rotor (Beckman Coulter) at 20,000 rpm for 1.5 h at 4 °C, and resuspended in hepatocyte defined medium (HDM; Corning). Primary human hepatocytes (Lonza) were transplanted into FNRG mice to create human liver chimeric mice as previously described [34]. Livers from highly humanized mice were harvested as described [35] and seeded on collagen-coated plates in HDM. Lentivirus transduction was performed over night in the presence of 8μg/ml polybrene followed by extensive washing. Transduced cells were harvested after 72 h and RNA was purified using RNeasy kit (Qiagen), including an on-column DNAse I treatment (Qiagen). cDNA was generated as described below.

### Ethics statement

All animal procedures were approved by Rockefeller University’s Institutional Animal Care and Use Committee under protocol number 18063-H. This protocol adheres to United States Department of Agriculture-Animal Welfare Act and Regulations, as well as to US Department of Health and Human Services and the National Institutes of Health-Public Health Service (PHS) Policy on Humane Care and Use of Laboratory Animals.

### RNA extraction and analysis

RNA was extracted using TRI Reagent (MRC) or the PerfectPure RNA Purification kit (5 PRIME). First-strand synthesis was performed using qScript cDNA synthesis kit (Quanta). qRT-PCR was performed using the LightCycler480 (Roche), with PerfeCta^®^ SYBR Green FastMix mix (Quanta). All qPCRs were normalized to 18S mRNA levels or RPS11 (PHH). The primer sequences are in the supplementary materials.

### Western blot

Lysates were prepared from cells using RIPA buffer [36] supplemented with Dithiothreitol (DTT) and protease and phosphatase inhibitors (Sigma), and subjected to SDS-PAGE.

#### Antibodies

Goat anti-R2 (Santa Cruz Biotechnology (SCB) N18 SC-10844), Mouse antiActin (Sigma A4700), Mouse anti-HA (Sigma), Rabbit anti-PSMA4 was a kind gift of C. Kahana, Weizmann Institute of Science, Rehovot, Israel), Rabbit anti-pChk1-ser345 (Cell Signalling Technology (CST) #2348), Rabbit anti-Chk1 (CST #2345), Rabbit anti-E2F-1 (SCB C-20 SC-193;), Rabbit anti-pATR-Thr1989 (Genetex GTX128145) and Mouse anti-ATR (SCB C-1 SC-515173). We used horseradish peroxidase–conjugated secondary antibody (Jackson), and enhanced chemiluminescence (ECL) detection using EZ-ECL (Biological Industries).

Quantification of western blot bands was done using ImageJ (version 1.51k) software. (http://imagej.nih.gov/ij).

### Phos-Tag gel

Phos-Tag™-AAL-107 was purchased from Wako Pure Chemical Industries, Ltd. (Japan), and was added at 40μM concentration to 6% SDS-PAGE gel.

### Constructs used for lentiviral transduction

Lenti-GFP and Lenti 1.3xHBV are based on the pHR’ vector [8]. Lenti-1.3xHBV and lenti-HBV-X-KO were cloned from pGEM3Z-1.3xHBV and-pGEM3Z-X-KO, which contains a stop codon in place of aa27KO as described [4]. Lenti-HBV-Core-KO was created by cloning the entire 1.3xHBV fragment from pGEM3Z-HBV-Core-GFP, in which Core ORF was replaced with GFP, as described in [37] into pHR vector, as was done for pHR-HBV. Lenti-HBV-S-KO was cloned similarly from pGEM3Z-HBV-S-GFP.

The constructs containing HBx ORF were clone into a pLenti4 Gateway vector (Invitrogen) and contained a HA-tag 5’ to the start of the sequence and the endogenous HBx 3’UTR, if not mentioned otherwise, with specific modifications of the HBX ORF as described.

For experiments done with/without CMV promoter, ERE and control sequences were cloned into a FUGW lentiviral vector (a gift from David Baltimore, Addgene plasmid # 14883) [38], with or without an upstream CMV promoter.

For expression of ERE under the U6 promoter, ERE was cloned into a pKLV vector (a gift from Hiroshi Ochiai, Addgene plasmid # 62348) [39].

### Statistical analysis

Error bars refer to standard error of the mean (SEM). A two-sided Student’s t-test was performed to assess significance.

Data analysis was preformed with a web-tool for plotting box plots http://shiny.chemgrid.org/boxplotr/ and Microsoft Excel.

## Acknowledgements

The Authors would like to thank Lior Handler and Nurit Papismadov for their help with the experiments.

## Supporting figure captions

**Fig S1. HBV activates RNR-R2 by activation of DDR pathway components in quiescent cells.** Upon HBV infection key enzymes in the DNA damage response pathway Chk1 and E2F1 are activated. This leads to transcription of the RNR-R2 gene, the formation of the RNR complex and dNTPs synthesis.

**Fig S2. HBx upregulates RNR-R2.** A) Quiescent HepG2 cells were transduced with Lenti-GFP (vector), Lenti-1.3xHBV or Lenti-HBx. HBx RNA levels were measured by qRT-PCR B) Cells were treated as in A. Shown is a representative western blot of RNR-R2 protein levels. Actin was used as a loading control. C) As in B, RNR-R2 mRNA levels were measured by qRT-PCR from three biological replicates.

**Fig S3: Pol ORF is not required for RNR-R2 upregulation.** A) A stop codon was introduced into the Lenti-HA-X construct, to prevent expression from the overlapping 3’ end of the Pol ORF. B) Relative RNR-R2 levels were measured by qRT-PCR from three biological replicates, where WT HA-X acts as positive control, and no-X-, a vector that contains the HA-tag and the 3’UTR sequences, but no HBx coding sequence, was used as a negative control.

**Fig S4. The endogenous 3’ UTR sequence is not required for RNR-R2 upregulation and neither is the 5’ flanking region.** A) The endogenous HBV 3’ UTR was removed from the HA-X-3’UTR construct, as depicted in the illustration. B) The resulting construct was transduced into quiescent HepG2 cells, and RNR-R2 induction was measured by qRT-PCR, as percentage of RNR-R2 induction by the original construct. C) Relative transduced RNA expression was measured by qRT-PCR, to verify similar amounts of RNA. D) The HA sequence upstream to the HBx coding region was removed, as depicted in the illustration, and the resulting construct was transduced into quiescent HepG2 cells. E) Relative RNR-R2 mRNA levels were measured by qRT-PCR. HA-HBx was used as positive control, no-X-, a vector that contains the HA-tag and the 3’UTR sequences, but no HBx coding sequence, was used as a negative control for RNR-R2 induction. F) Transduced RNA expression was measured by qRT-PCR, with primers from the shared 3’UTR region.

**Fig S5. ATM inhibition does not affect ERE-mediated RNR-R2 induction.** A) Noncycling HepG2 cells expressing either ERE or the control RNA were treated with 2μM Caffeine (ATM/ATR inhibitor) or 1μM KU55933 (ATM inhibitor) for 24h. Relative RNR-R2 mRNA levels were measured by qRT-PCR. B) Transduced RNA levels were measured by qRT-PCR to validate they remain unchanged following treatment with the indicated inhibitors. C) To validate the inhibitors, HepG2 cells were treated with the indicated inhibitors, irradiated with X-ray (5Gy), to induce DNA damage response, and harvested after 24h. Relative p21 mRNA levels were measured as an indication for DDR inhibition.

